# The contribution of lincRNAs at the interface between cell cycle regulation and cell state maintenance

**DOI:** 10.1101/848333

**Authors:** Adriano Biasini, Adam Alexander Thil Smith, Baroj Abdulkarim, Jennifer Yihong Tan, Maria Ferreira da Silva, Ana Claudia Marques

## Abstract

Cell cycle progression requires dynamic and tightly-regulated transitions between well-defined cell cycle stages. These transitions are controlled by the interplay of established cell cycle regulators. Changes in the activity of these regulators are thought to underpin differences in cell cycle kinetics between distinct cell types. Here, we investigate whether cell type-specific long intergenic noncoding RNAs (lincRNAs) contribute to embryonic stem cell adaptations, which have been shown to be essential for the maintenance of embryonic stem cell state.

We used single cell RNA-sequencing data of mouse embryonic stem cells (mESC) staged as G1, S, or G2/M to identify genes differentially expressed between these phases. We found differentially expressed lincRNAs to be enriched amongst cell cycle regulated genes. These cell cycle associated lincRNAs (CC-lincRNAs) are co-expressed with protein-coding genes with established roles in cell cycle progression. Interestingly, 70% of CC-lincRNAs are differentially expressed between G1 and S, suggesting they may contribute to the maintenance of the short G1 phase that characterizes the embryonic stem cell cycle. Consistent with this hypothesis, the promoters of CC-lincRNAs are enriched in pluripotency transcription factor binding sites, and their transcripts are frequently co-regulated with genes involved in the maintenance of pluripotency. We tested the impact of 2 CC-lincRNA candidates and show that modulation of their expression is associated with impaired cell cycle progression, further underlining the contribution of mESC-specific lincRNAs to cell cycle modulation in these cells.

## INTRODUCTION

The cell cycle is a dynamic sequence of events leading from one parent cell to two daughter cells. This requires replication of chromosomes during the Synthesis phase (S phase) and their segregation into two daughter cells during Mitosis (M phase). M and S phases are separated by 2 Gap phases, G1 and G2, that act as checkpoints to prevent cell division without genome replication and aberrant polyploidy [1]. Progression through cell cycle is regulated by cell cycle stage-specific activation and repression of numerous proteins including Cyclin Dependent Kinases (CDKs) and proteins from the Cyclin family [2]. In most somatic cells, the oscillatory expression or activity of distinct Cyclin-Cdk complexes allows the activation and repression of cell cycle regulators and promotes cell-cycle transitions [2]. One key regulator of this process is the retinoblastoma protein (RB) that controls G1 and prevents entry into S phase. Upon entrance in G1, RB is unphosphorylated (active) and blocks the expression of genes required for G1/S transition. During G1, RB is phosphorylated and becomes inactive, allowing cells to progress to S phase [3]. In embryonic stem cells (ESCs), RB is hyperphosphorylated, resulting in suppression of the G1-S checkpoint and thereby rapid shuttling between DNA synthesis and mitosis, decreasing the average duration of the ESC cell cycle [reviewed in [4]].

These significant adaptations in the ESC cell cycle are important for the maintenance of the embryonic stem cell state and cell fate decisions, as highlighted by the partial overlap between the gene regulatory networks that control the two processes [5]. For example, both Oct4 and Nanog, two core pluripotency factors, control genes involved in cell cycle regulation: in mouse ESCs (mESCs), Oct4 represses the expression of *p21*, a Cyclin-dependent kinase inhibitor that is expressed in somatic cells but not in embryonic stem cells [6]; *NANOG*, whose expression is cell cycle-regulated, controls S-phase entry by regulating the expression of *Cdc25C* and *CDK6* in human ESCs (hESCs) [7]. While the association between cell cycle dynamics and cell state is well established, the molecular mechanisms underlying this connection remain uncharacterized [5].

In addition to proteins, noncoding RNAs, including long intergenic noncoding RNAs (lincRNAs), have also been shown to contribute to cell cycle progression [8]. An example of this is MALAT1, a lincRNA that is frequently upregulated in multiple human cancers [9]. In human fibroblasts, depletion of MALAT1 leads to decreased expression of several genes involved in cell cycle progression and results in G1 arrest. MALAT1 is also involved in splicing of *B-Myb*, a gene involved in the transcriptional regulation of several mitotic proteins [10]. More recently, the cohesion regulator long noncoding RNA (CONCR) has been found to be necessary for cell cycle progression and DNA replication. CONCR expression is activated by the transcription factor MYC and is upregulated in multiple cancer types [11]. Silencing of CONCR leads to a significant decrease in DNA synthesis. At the molecular level, CONCR physically interacts with DDEA/H-boy helicase 11, which ensures the proper separation of sister chromatids during the cell division process. The absence of CONCR leads to the loss of sister chromatid cohesion and affects metaphase [11]. Finally, lincRNA expression is often dysregulated in cancer and the characterization of subsets of cancer-associated lincRNAs highlights their potential roles as cell cycle progression modulators [12, 13].

LincRNAs are also part of the network controlling stem cell fate maintenance and differentiation [14]. Because lincRNA expression is often tissue-specific [15, 16], in contrast to proteins, we hypothesized they can support cell type-specific activity of ubiquitously-expressed genes and act at the intersection of cell cycle and mESC cell state regulation. The ability of tissues specific noncoding RNAs to modulate the activity of ubiquitously expressed gene was already exemplified by lncSCA7. This lincRNA regulates ATXN7 levels in retinal and cerebellar neurons and contributes to specific degeneration of this cells in SCA7 patients [17]. Additionally, the relative short half-lives of lincRNAs [18, 19] further underscores their potential as modulators of temporally resolved processes such as cell cycle progression.

Here, we investigate the contributions of lincRNAs to embryonic stem cell cycle adaptations.

## RESULTS

### LincRNA expression is often cell-cycle regulated

We used publicly-available single-cell RNA-sequencing (scRNAseq) data for 279 mouse embryonic stem cells with known cell cycle stage [20] to assess the extent of cell cycle-regulated lincRNA expression in mouse embryonic stem cells (mESCs). We estimated the expression of protein-coding transcripts and lincRNAs in each of these cells. After excluding cells and genes that failed quality control (Supplementary Figure 1) we identified 10 487 genes, including 781 lincRNAs, whose expression can be robustly detected in 246 cells. As previously shown [20], gene expression patterns in this dataset reflect the cell cycle stages of the individual cells (Figure 1A), supporting its use to identify transcripts whose expression are cell-cycle dependent. Using DEseq2, we identified 638 genes (6.1%) whose expression is significantly different between at least 2 cell cycle stages (Supplementary table 1). The proportion of differentially expressed lincRNAs (n=70, 8.96%) is significantly higher than found for protein-coding genes (n=501, 5.51%) (proportions test p-value < 0.05, Figure 1B, Supplementary Table 1), indicating that lincRNA expression is more dynamic throughout the mESC cell cycle than is the expression of protein-coding genes.

**Figure 1:**
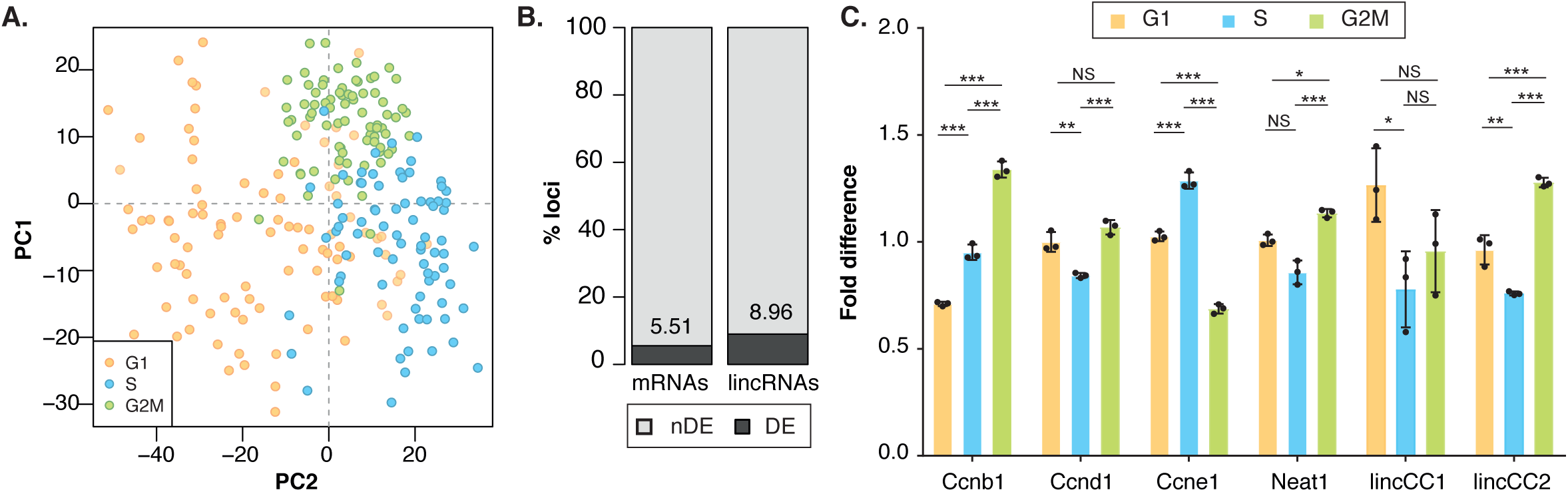
lincRNA expression is often cell cycle regulated. **A)** Principal component analysis of gene expression for all loci passing the technical noise filter. The first 2 principal components (PC1 on the x-axis and PC2 on the y-axis) together explain 5.15 % of the total variability, and separate cells in different cell cycle stages (G1: orange, S: blue; G2/M: green). **B)** Percentage of protein-coding genes (left) and lincRNAs (right) that are differentially expressed (dark grey, numbers indicate the percentage) between at least 2 cell cycle stages. **C)** Fold difference in normalized expression (relative to G1) of known cell cycle regulated genes (Ccnb1, Ccnd1, Ccne1, Neat1) and 2 differentially expressed novel lincRNAs (lincCC1 and lincCC2) in mESCs at different cell cycle stages (G1: orange, S: blue, G2/M: green). Significance (two-tailed unpaired t-test p-value) between cell cycle stage expression is indicated as: NS p> 0.05, * p<0.05, ** p<0.01, *** p<0.001.

The median expression of mRNAs is roughly 14 times higher than that of lincRNAs (Supplementary Figure 2A) and steady state abundance can impact the ability to detect differential gene expression, as highlighted by the significantly higher expression (two-tailed Wilcoxon test, p-value< 2×10^−16^) of genes identified as differentially expressed (Supplementary Figure 2A). To assess whether lincRNAs’ relatively low expression impacts our differential gene expression analysis results, we repeated the analysis by re-quantifying protein-coding gene expression using a subset (1/14) of randomly sampled reads from each library. At this sequencing depth, the median mRNA expression is comparable to that of lincRNAs in the full data (Supplementary Figure 2B). We repeated expression quantification, quality control, normalization, filtering and differential gene expression analyses using these randomly sampled reads and found that only 60% of the mRNAs found to be differentially expressed using the full data are also differentially expressed when using the down-sampled libraries. This result indicates that the higher proportion of differentially expressed lincRNAs is likely an underestimate due to the limited power to measure their expression using single cell RNA sequencing data.

**Figure 2:**
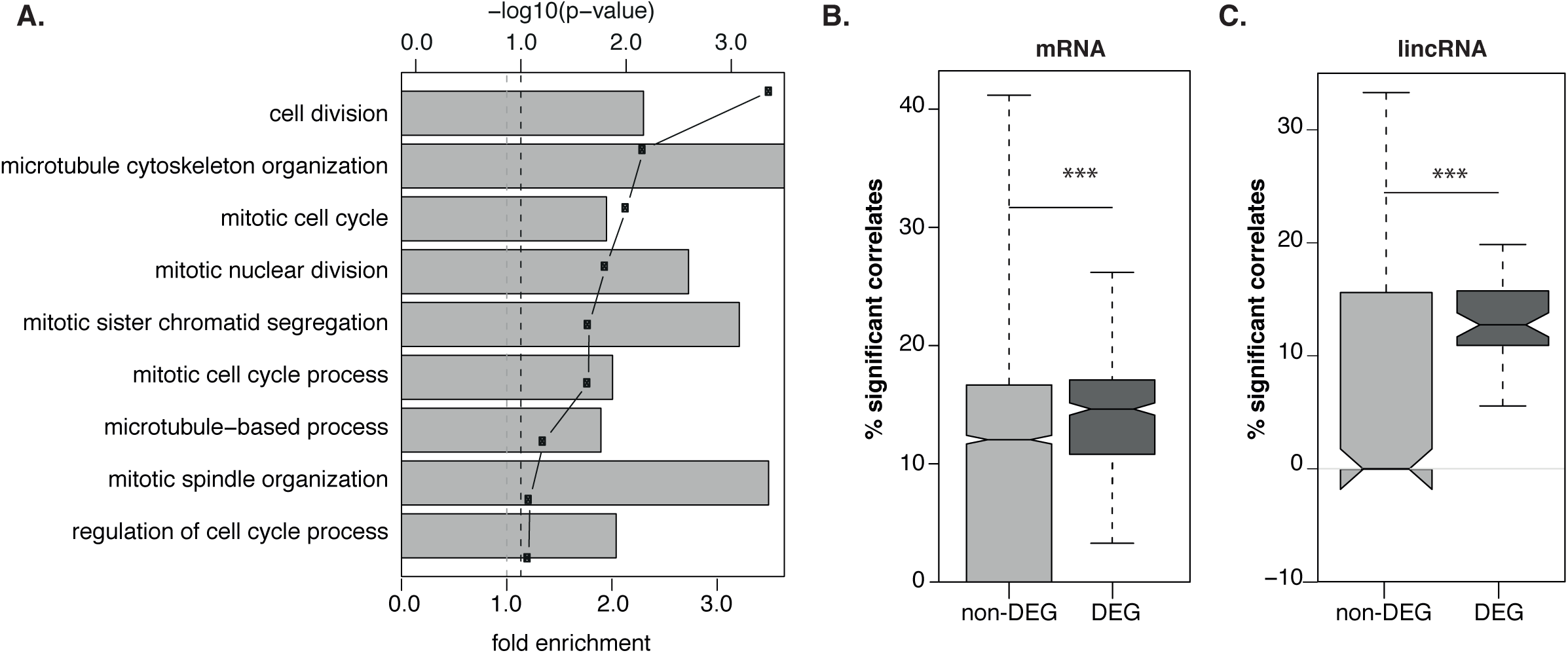
Genes differentially expressed between cell cycle stages have cell cycle related functions. **A)** Results of the GO term enrichment analysis for cell cycle stage differentially expressed genes. Grey bars indicate the fold enrichment, and points the associated −log10 of the p-value. Dashed line indicates the 10% FDR cutoff. Distribution of the percentage of genes cell cycle regulators that are significantly correlated in expression with **B)** protein-coding genes and **C)** lincRNAs. Differentially or non-differentially expressed genes from each biotope are represented in dark and light grey respectively. Significance of the distribution comparison (two-tailed Mann-Whitney-Wilcoxon rank-sum test) are indicated as: *** p-value < 0.001.

To validate our *in silico* differential gene expression predictions we used DNA content to sort mESC cells into G1, S and G2/M stages and measured the cell cycle stage expression of 12 differentially expressed lincRNAs, including Neat1, and 3 mESC cell cycle protein-coding genes (*Ccnb1, Ccnd1* and *Ccne1*) by quantitative PCR (Figure 1C, Supplementary Figure 3). The cell cycle protein coding gene and lincRNA expression patterns measured by qPCR were generally consistent with what was estimated using scRNA sequencing data. The results of this analysis support that despite the relatively low expression of lincRNAs that complicates the accurate estimation of their expression, our differential expression predictions are generally robust.

**Figure 3:**
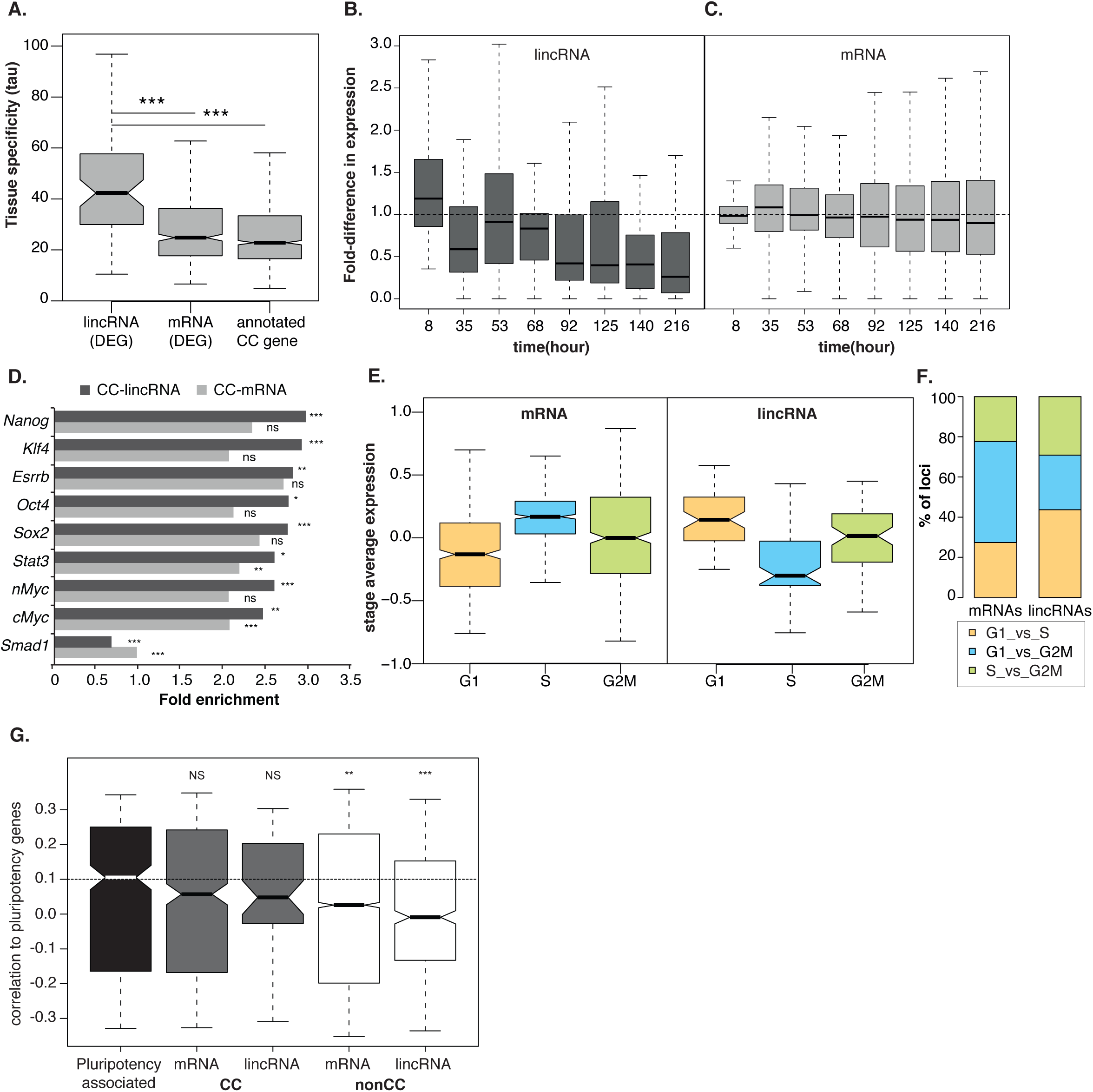
CC-lincRNAs are associated with stem cell cycle adaptations. **A)** Distribution of tissue specificity (measured as Tau) for CC-lincRNAs, CC-PCGs, and annotated cell cycle genes. Distribution of the fold difference in expression of **B)** lincRNAs and **C)** protein-coding genes during a 216-hour neuronal commitment of mESCs. Fold difference in expression is relative to the median fold difference at time=8 hours. **D)** Fold enrichment in binding (bars, x axis) of pluripotency transcription factors (y-axis) for CC-PCGs (dark grey) and CC-mRNAs (light grey). **E)** Cell cycle stage-average (G1: orange, S: blue, G2/M: green) of the expression Z-scores of CC-PCGs (left panel) and CC-lincRNAs (right panel). **F)** Percentage of protein-coding genes (left) and lincRNAs (right) differentially expressed between G1 vs S (orange), G1 vs G2M (blue) or S vs G2/M (green). **G)** Distribution of the correlations between pairs of pluripotency-associated genes (black) or known pluripotency genes, and differentially expressed (CC, grey) or non-differentially expressed (nonCC, white) protein-coding genes and lincRNAs. Significance (two-tailed Mann-Whitney-Wilcoxon rank-sum test) for the comparison between pluripotency-associated genes and the different gene classes is indicated on top of the boxplot as: NS p> 0.05, ** p<0.01, *** p<0.00.

### Differentially expressed lincRNAs contribute to the regulation of cell cycle progression

Consistent with the role of differentially expressed protein-coding genes in the regulation of cell cycle progression, these genes are significantly enriched (hypergeometric test adjusted p-value < 0.001) in annotations with gene ontology terms such as “cell division” and other related cell-cycle processes (Figure 2A). Furthermore, 8.8% (44 of 501) of differentially expressed protein-coding loci are orthologous to genes previously shown to have a periodic expression throughout the human cell cycle [21], a significant 3.1-fold enrichment relative to all mESC expressed protein-coding loci (214, hypergeometric test p-value < 0.05). These results support that genes differentially expressed between cell cycle stages are indeed enriched in cell cycle regulators.

The enrichment of cell-cycle related genes amongst those differentially expressed between cell-cycle stages suggests that some of the lincRNAs identified here as differentially expressed might also contribute to mESC cell cycle regulation. Most lincRNAs differentially expressed between cell cycle stages were annotated *de novo* (53 *de novo* lincRNAs) using mESC RNA sequencing data [22]. We searched the literature for functions of the remaining 17 differentially expressed lincRNAs annotated by ENSEMBL. Of these, only 4 have been previously characterized: Neat1, Malat1, the host transcript for snord93, and miRNA-24 pri-miRNA transcript. Malat1 is a cancer-associated lincRNA and has been previously implicated in cell cycle progression. For example, Malat1 is highly expressed during the S and M phases of human Fibroblasts where it controls cell cycle-related processes [10]. Neat1, whose knockdown was recently shown to impair S phase transition [23] has also been frequently associated with cancer and cell cycle progression [24]. Relatively little is known about snord93 but changes in this RNA’s levels impact cell proliferation [25], which would be consistent with a role in cell cycle progression. Finally, miRNA-24 post-transcriptionally regulates MYC and E2F2, two known cell cycle genes, and changes in its levels impair G1 transition [26]. In conclusion, all of the 4 annotated and characterized lincRNAs we identified as being differentially expressed either have established roles or have been associated with cell cycle progression.

Given the paucity of functional annotations for lincRNAs, and to assess broadly the contributions of differentially expressed lincRNAs to the cell cycle, we reasoned that genes that functionally participate in cell cycle regulation should genetically interact with known cell cycle regulators. To first validate this idea, we considered cell cycle phase differentially expressed protein-coding genes not annotated as cell cycle regulators, and found their expression is more often significantly (p < 0.001, Wilcoxon rank sum test) correlated with the expression of annotated cell-cycle genes than non-differentially expressed protein-coding loci (Figure 2B). Similarly, we found that the expression of differentially expressed lincRNAs is also significantly more often correlated with the levels of cell cycle genes than non-differentially expressed lincRNAs (Figure 2C, p < 0.001 Wilcoxon rank sum test), supporting their contributions to mESC cell cycle progression. Hereafter, we refer to differentially expressed lincRNAs and protein-coding genes as cell-cycle associated lincRNAs and protein coding genes, or CC-lincRNAs and CC-PCGs, respectively.

### Cell cycle lincRNAs are associated with stem cell cycle adaptations

The cell cycle is a ubiquitous process, yet most lincRNAs are expressed in cell-type specific manner [27]. Do the CC-lincRNAs identified here contribute to cell cycle regulation ubiquitously, or are their functions restricted to mESCs? To gain initial insights into this question we used publicly available transcriptome-wide data [28] to estimate the tissue specificity of lincRNAs and protein coding genes across 29 adult and developing mouse tissue and cell lines. We estimated Tau, a measure of tissue specificity, for CC-lincRNAs, CC-PCGs and known cell cycle regulators. Tau varies between 0% for “ubiquitously expressed” to 100% for “tissue-specifically expressed” genes [29]. As expected, CC-PCGs, as well as established cell-cycle genes, are broadly expressed (median tau=24.8%). In contrast, CC-lincRNAs are significantly more tissue specific (median tau=42%, Wilcox rank sum test p-value < 0.05, Figure 3A) than their protein-coding counterparts, and are as tissue-specific as other mESC expressed lincRNAs (data not shown).

To further test if CC-lincRNA expression is often restricted to mESCs, we investigated changes in their transcript abundance upon neuronal differentiation of mESCs. *In vitro* neural differentiation is a well-defined and highly efficient process (>80% of differentiated cells). We took advantage of publicly-available transcriptome-wide expression for a time-course of neuronal commitment of mESC [30] to investigate the changes in noncoding and coding gene expression. Consistent with their tissue-specific expression (Figure 3A), CC-lincRNA expression decreases rapidly upon differentiation (Figure 3B), supporting their contribution to cell cycle regulation being restricted to mESCs. In contrast, CC-PCGs are expressed at similar levels throughout neuronal commitment (Figure 3C).

The transcriptional network controlling mESC cell state and function is regulated by a set of mESC core transcription factors [31]. The short G1 phase that characterizes the embryonic stem cell cycle, which is critical to ensure maintenance of pluripotency, is in part orchestrated by stem cell-specific factors [32]. We took advantage of publicly available data ChIP-seq data for pluripotency transcription factors in mESCs [33], to assess the extent of these factors’ binding of to cell-cycle regulated lincRNA promoters. We found that CC-lincRNA promoters are enriched in mESC core transcription factors (TFs) supporting their role in the network underlying mESC cell state. Specifically, promoters of CC-lincRNAs are significantly enriched, relative to all expressed lincRNAs, in the binding of pluripotency transcription factors, including Nanog, Oct4 and Sox2 (FDR<0.05, permutation test) (Figure 3D) We found no significant enrichement by most pluripotency TFs at the promotes of CC-PCGs (Figure 3D).

To assess what aspect of mESC cell cycle progression might be more often modulated by CC-lincRNAs, we investigated their relative expression across different cell cycle phases. Consistent with previous observations that mRNA expression peaks at G2/M phase in human cells [34, 35] we found that CC-protein-coding genes were highly expressed in S and G2/M (Figure 3E) and were often differentially expressed between G1 *vs* G2/M (Figure 3F). In contrast, the levels of CC-lincRNAs were higher in G1 relative to all other cell cycle stages (Figure 3E) and most were differentially expressed between G1 *versus* S phase (Figure 3F). In total, 70% of all CC-lincRNAs are differentially expressed between G1 and another cell cycle stage.

Given the significantly elevated expression of CC-lincRNAs in G1, their tissue-specific expression and evidence that their transcription is regulated by stem cell-specific transcription factors, we hypothesized that CC-lincRNAs may contribute to the interplay between cell cycle and maintenance of pluripotency in mESCs. If CC-lincRNAs participate in the network controlling maintenance of mESC cell state, they should be co-expressed with genes involved in maintenance of pluripotency. To test this hypothesis, we investigated the correlation, during mESC neuronal commitment, between CC-genes and genes implicated in maintenance of pluripotency [36]. First, we found that, consistent with the interplay between cell cycle control and maintenance of mESC cell state, the median pairwise correlation between CC-PCGs and pluripotency genes (Spearman’s r=0.06) is similar to what is found for pairs of genes implicated in pluripotency (r=0.10, two-tailed Wilcoxon test p-value=0.3, Figure 3G). Consistent with our hypothesis that CC-lincRNAs contribute to maintenance of mESC cell state, we found that the extent of their association with pluripotency genes (r=0.05) is also similar to what we estimate for pairs of genes implicated in pluripotency (two-tailed Wilcoxon test p-value=0.7, Figure 3G). For comparison, we also estimated the strength of the association with genes involved in pluripotency for non-cell cycle differentially expressed mESC coding (r=0.03) and non-coding (r=-0.01) genes and found this to be significantly lower than that of CC-lincRNAs or CC-mRNAs (two-tailed Wilcoxon test p-value<0.05, Figure 3G). Similar results are obtained when considering only significant correlations (correlation test p<0.05) between pluripotency gene and coding or noncoding transcripts.

These results suggest that CC-lincRNAs modulate specific aspects of mESC cell cycle and may contribute to the regulation of stem cell cycle adaptions.

### Candidate CC-lincRNA analysis supports lincRNA roles in cell cycle regulation

Next, we aimed to experimentally validate the contributions of previously uncharacterized CC-lincRNAs to mESC cell cycle regulation. Considering that relatively low transcript abundance may limit individual lincRNA’s ability to regulate cell cycle progression, we selected one modestly expressed CC-lincRNA (XLOC_018328, hereafter lincCC1, Supplementary Figure 4A), and one lincRNA that is relatively highly expressed (Gm26853, hereafter lincCC2, Supplementary Figure 4B) in mESCs (Figure 4A). We employed CRISPR activation (CRISPRa) to increase endogenous lincRNAs’ transcription. For each lincRNA, we designed 2 guide RNAs (gRNA) to target VP160 fused dead-Cas9 (dCas9-VP160) [37] to the vicinity of lincRNA promoters. As a control, we designed a non-targeting scrambled gRNA. We transiently co-transfected each of the gRNA-expressing constructs with a dCas9-VP160 expressing vector. Seventy-two hours post-transfection we observed an average 2-fold upregulation (2-tailed unpaired t-test p-value < 0.05) of candidate lincRNAs expression (Figure 4B-C) in mESCs treated with targeting gRNA, relative to control.

**Figure 4:**
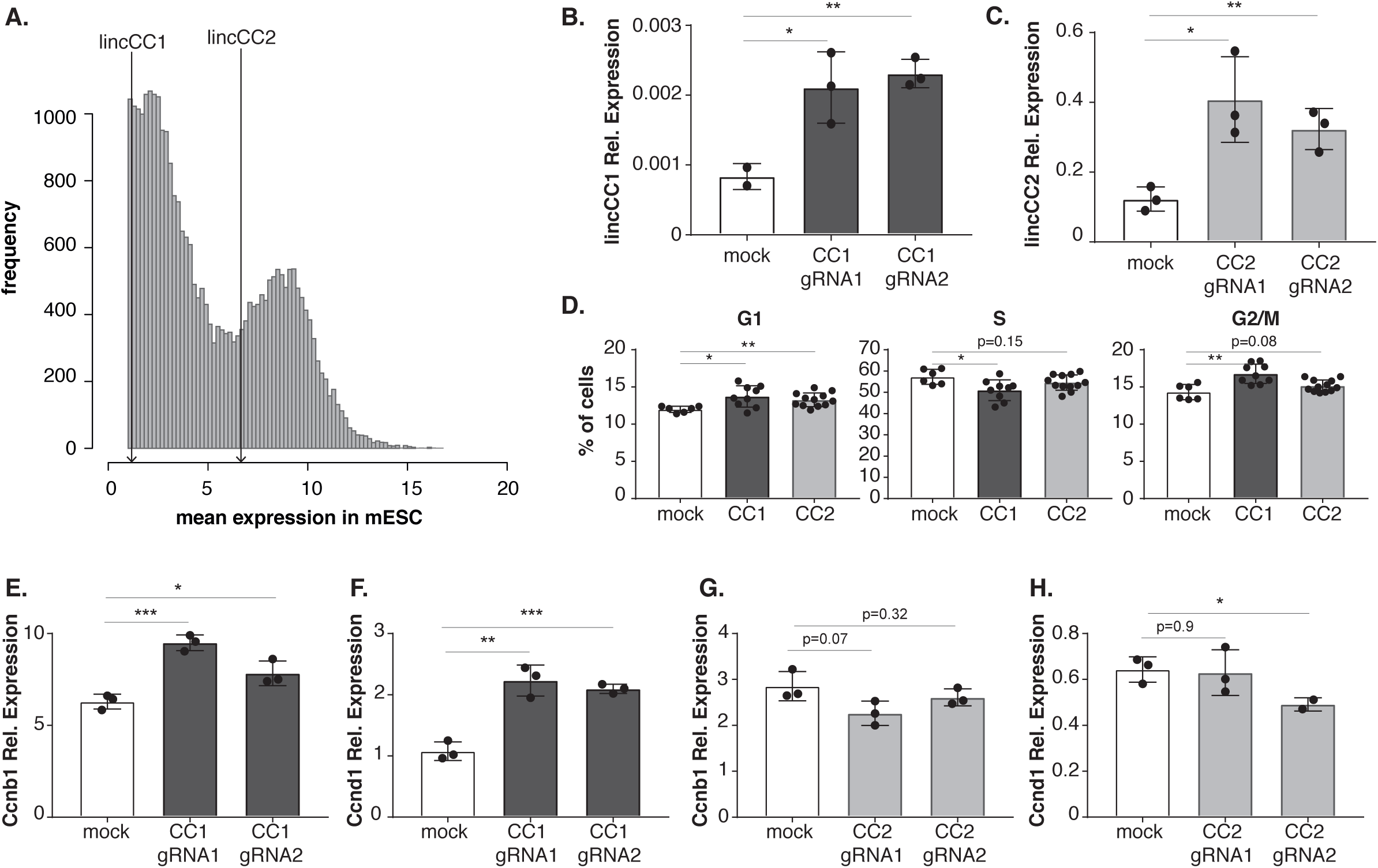
Candidate lincRNA overexpression impacts cell cycle progression. **A)** Histogram of the mean cell expression of all considered loci in mESCs. Lines indicate the expression of lincCC1 and lincCC2. Relative **B)** lincCC1 and **C)** lincCC2 expression 72 h after co-transfection of Cas9-VP160 with scramble-, gRNA1- or gRNA2-expressing constructs. **D**) Percentage of mESCs at G1, S and G2/M stages of the cell cycle 72 h post-transfection of mESCs with scramble (white), lincCC1 (dark grey) or lincCC2 (light grey) targeting gRNAs. Relative expression of *Ccnb1* (**E-G**) and *Ccnd1* **(F-H)** 72 h post-transfection of mESCs with scramble (white), lincCC1 (dark grey) or lncCC2 (light grey) targeting gRNAs. Significance (two-tailed unpaired t-test p-value) of comparisons is indicated as: NS p> 0.05, * p<0.05, ** p<0.01, *** p<0.001.

To assess the impact of changes in candidate lincRNA expression upon cell cycle progression in mESC, we determined the proportions of cells in G1, S and G2/M phases using Fluorescence Activated Flow Cytometry (FACS). We used EdU incorporation and DNA content to identify the proportion of single cells in each cell cycle phase following CC-lincRNA overexpression(Supplementary Figure 4C). We found that, consistently with the proposed role of lincCC1 and lincCC2 in the modulation of cell cycle progression, upregulation of these lincRNAs is associated with significant changes in the proportion of cells within different cell cycle stages. LincCC1 upregulation is associated with a significant increase in the number of cells in G2/M and small, yet significant changes in the proportion of cells in G1 and S phases (Figure 4D). LincCC2 mostly affects the proportion of cells in G1. We have also investigated changes in cell cycle markers, specifically *Ccnb1* and *Ccnd1*, whose expression in mESC is highest in G2/M and lowest in G1. As expected, we found that lincCC1 upregulation is associated with a significant increase in the two cell cycle markers (Figure 4E-F). Increased expression of lincCC2 moderately decreased the expression of both markers which is consistent with the increase in the number of cells in G1 (Figure 4G-H).

These observations are consistent with the role of these CC-lincRNAs in the regulation of cell cycle progression.

## CONCLUSION

The cell cycle is central to the development of a multicellular organism from a single cell zygote and is required for cell renewal. Dysregulation of this process can cause disease, most notably cancer. Progression through the cell cycle requires dynamic and tightly-regulated transitions between well-defined cell-cycle stages, which are controlled by changes in cell cycle regulators’ activity. While control of cell cycle progression has been primarily assigned to protein-coding genes, several lincRNAs have recently been shown to contribute to this process [8]. For example, a recent siRNA screen in HeLa cells revealed that knockdown of 26 lincRNAs, whose expression is deregulated in cancer, impacts different aspects of cell cycle progression [13]. The role of lincRNAs in this process is further supported by a more recent comprehensive high-content RNAi screen in the same cells [38]. In parallel, characterization of individual lincRNAs has also illustrated how these noncoding RNAs contribute to the regulation of different aspects of cell cycle [8].

In contrast to most cell cycle protein-coding regulators, lincRNA expression is often restricted in time and space, as supported by the analysis of their expression across adult and embryonic tissues and in single cells [for example [15, 16, 27]]. This suggests that most lincRNA-encoded functions are tissue-specific. While lincRNAs have been previously implicated in the regulation of cell cycle progression [10, 11, 23], their role, if any, in cell type-specific modulations of this central biological process remains understudied. Here, we used mouse embryonic stem cell (mESC) cycle, which is characterized by a truncated G1 phase, as a model system to assess the contributions of lincRNAs expressed in mESCs to the modulation of this stem cell adaptation.

To identify lincRNAs that are putative cell cycle regulators, we reasoned that for noncoding genes the functional moiety is the transcript, thus differential noncoding gene expression between cell cycle stages would result in differential activity and enrich for lincRNAs with roles in cell cycle regulation. Consistent with this hypothesis, our genome-wide analyses and experimental validations support the role of lincRNAs differentially expressed between mESC cell cycle stages as modulators of cell cycle progression, likely through interaction with other cell cycle regulators.

Interestingly and relative to protein-coding genes, whose activity during cell cycle is often modulated by post-translational modifications, lincRNAs are enriched among differentially expressed transcripts, supporting that in mESCs their expression is more frequently dynamic throughout cell cycle progression. The fraction of cell cycle-regulated lincRNAs is similar to a previous estimate done in HeLa cells using bulk RNA sequencing (∼9%,[39]). However, the dynamics of lincRNA expression throughout the cell cycle differs between cell types, as in contrast with this earlier study in HeLa [39] that revealed no preferential cell stage specific expression in mESCs, we found most lincRNA are highly expressed in G1 and differentially expressed between G1 and S phase. Given the critical importance of the G1-S transition in maintenance of embryonic stem cell state [32], we hypothesize that a subset of lincRNAs with roles in mESC cell cycle progression also have roles in maintenance of stem cell state. Supporting this hypothesis is the observation that these lincRNAs’ expression is often restricted to pluripotent cells, and rapidly decreases upon exit from pluripotency and entry into neural commitment. Furthermore the expression of these lincRNAs is often regulated by core pluripotency transcription factors, including Nanog, Sox2 and Oct4. Finally, we provide preliminary evidence that cell cycle lincRNAs are part of the network underlying pluripotency.

While further work is now required to disentangle how individual lincRNAs may contribute to cell cycle progression in mESCs, our results suggest that tissue-specific regulation by lincRNAs contributes to cell-type specific adaptations of ubiquitous processes.

## METHODS

### Processing of cell cycle staged single cell RNA sequencing data

Mouse genome sequence and annotation files were downloaded from ENSEMBL (GRCm38.82 i.e. mm10). LincRNAs expressed in mESCs [17] were added to the transcript annotations using custom Python scripts in order to appropriately deal with novel transcripts overlapping existing loci, leading to a total of 55,596 loci. The FASTA sequences of the 96 ERCC spike-ins used in the experiment were downloaded from UCSC GoldenPath. We used CGAT ([40], v 0.2.3) to extract transcript sequences longer than 200 nucleotides, and Kallisto ([41], v 042.4, default parameters) to build a transcriptome index that included 114,842 transcripts across 49,285 loci and ERCCs. The FASTQ files for staged mESC single-cell RNA-sequencing were downloaded from ArrayExpress (accession number E-MTAB-2805). For each cell, we estimated transcript expression using Kallisto (v. 0.42.4) [41] without bootstraps. Hit counts were imported into R and summarized to the gene level using tximport [42]. After removing genes without any hits in any cells, only 38,016 genes remained.

Basic cell-level quality control was performed as previously described [20]. Briefly, we excluded cells: 1) with a cell-wise gene detection rate below 20%; 2) cells with <1.6M total hits; 3) cells with proportions of hits to ERCCs lower than 15% or higher than 65%; 4) cells with a proportion of hits to mitochondrial genes lower than 1% or higher than 10%; and 5) cells with an estimated cell size higher or lower than 2 median absolute deviations from the stage-median cell size. In total, after these filtering steps, 94/95 G1 cells, 76/88 S cells, and 74/96 G2/M cells were retained for further analysis.

We then performed gene-level quality control. We considered only genes that had at least one hit count in at least 4% of samples. Hit counts were normalized using the scran package (v1.0.4, [43]). Size factors were calculated using pools of samples of sizes 10 to 35 cells from each stage. We used scLVM [20, 44] to filter out genes whose total variance is not significantly higher than that expected from pure technical variability (fitTechnicalNoise parameters: mincv2=0.01, quan=0.10; getVariableGenes parameters: threshold = 0.10, minBiolDisp = 0.30), and subsequently also removed ERCCs & mitochondrial genes. After this final filtering step, 10,487 genes were kept.

Except for differential expression analysis, where scran-normalised counts were used, all subsequent analyses were performed on scran-normalised shifted log10 counts (sl10 = log10(normalised counts + 1)).

### Processing of bulk RNA sequencing for a time-course of mESC to neurons differentiation

We used STAR [45] (v 2.5.3a) to map bulk RNA sequencing data for a 216-hour time course of mESC differentiation into neurons [30] to the Mouse transcriptome. Transcript quantification was performed using RSEM [46] (v 1.3.0) and for each time point we considered the average of expression across time points. We estimated the Spearman correlation between genes of interest and pluripotency associated genes [36] using R.

### Differential expression analysis

Kallisto pseudo-count data was imported into R (v 3.3.1) and prepared for DESeq2 (v 1.12.4) [47] using the *tximport* [42] package (v 1.0.3).

Principal Components Analysis (PCA) was applied using R package FactoMineR [48] to an expression matrix consisting of all cells passing QC in rows, and all genes passing QC in columns. The cell cycle signal typically appeared most strongly across the first two principal components, with the third component correlating strongly with the gene dropout rate (not shown). Differentially expressed genes (FDR< 5%, log2 fold change > 0.1) were called between pairs of cell cycle stages using DESeq2 [47, 49] using appropriate contrasts.

In order to assess whether the overall lower expression of lincRNAs affected our ability to call them as DE, we performed a down-sampling experiment to bring mRNA expression down to levels comparable to lincRNAs. We first estimated the average scaling factor between mRNA and lincRNA expression in the original data. We estimated the difference between the median Transcripts Per Million reads (TPM) in quality-controlled cells between lincRNAs and mRNAs to be 14.196, i.e. on average mRNAs are 14x more highly expressed than lincRNAs. We randomly sub-sampled the original FASTQ files to 1/14th of their original size (i.e. using a factor of 0.0714) using seqtk (v 2015.10.15). We quantified transcript expression using Kallisto and filtered cells and genes as described above prior to differential gene expression analysis. For the sub-sampled data, the number of tested genes was similar to the full data (9 923 vs 10 391). We then counted the number of mRNA genes called as DE in the full analysis that passed the quality control filters for the sub-sampled data (388), and the number of those called as DE in the sub-sampled analysis (233, i.e. 60.1%).

### Co-expression analysis

We considered a gene to be a cell cycle regulator if: 1) was annotated as a cell cycle gene [20]; 2) it was annotated with a GO term containing the string “cell cycle” or 3) had a Human ortholog annotated in CycleBase [21]. Pluripotency-associated genes were extracted from the ESCAPE database [36].

Co-expression analysis with cell cycle genes was performed using the single cell RNA sequencing data with staged mESCs [20]. We considered all loci expressed (normalized shifted log10>0.14) in at least 50 cells (out of a possible 244). The expression cut-off was defined based on inspection of the shifted log10 expression values. We estimated the Spearman correlation between all these loci and considered a gene pair to be significantly correlated if the absolute Spearman correlation between their expression was > 0.10 and the correlation test p-value was < 0.10. The correlation between any given pair was only calculated in cells where both genes were expressed. Correlations for gene pairs where there were less than 20 such value pairs were set to 0.

Co-expression analysis with pluripotency genes was performed using the time course of mESC to neurons differentiation data, as described above.

### Enrichment analyses

Enrichment analyses for Gene Ontology Biological Process, KEGG and REAC terms in *Mus musculus* were carried out using R package “gProfileR” (v. 0.6.1) [50]. Terms were considered to be significantly enriched if the associated corrected p-value was below 10% relative to the specified background of mESC expressed genes.

Enrichment in binding of mESC transcription factors (TFs) [33] at CC-lincRNA promoters (defined as +/-1 kb from their annotated TSS) was estimated using the Genome Association Tester (GAT) [40]. GAT tests whether the enrichment of TF binding at CC-lincRNA promoters is different from what would be expected based on 10,000 random samplings (with replacement) of intergenic segments with the same length and GC content as the CC-lincRNA promoters. The gat-compare tool was used to test the significance of the enrichment of TF binding at CC-lincRNA promoters relative to that at non CC-lincRNA promoters. Core TF binding sites were obtained from [33].

### Loci tissue specificity metrics in Mouse

We manually selected 150 Mouse paired-end total RNA bulk RNA-sequencing datasets from the ENCODE project [51], covering a range of tissues, sexes and ages, and added 3 publicly-available Mouse embryonic stem cell datasets [22]. We defined a “tissue.simple” grouping of samples, based on a high-level description of the tissue of origin (e.g. central nervous system, or heart) and of the developmental stage (adult or embryonic). Gene expression was estimated using Kallisto and imported into R using tximport as described above. Library size normalization was performed with DESeq2, and counts were transformed into shifted log10 normalised values. Values below a background level of 0.1 were set to 0. This cut-off was obtained based on the distribution of shifted log10 expression values. Genes with no expression values above 0 in any sample were discarded. Expression values were averaged across technical and biological replicates using the median. Tau was calculated as described in [29], briefly as Tau = sum(1-(v/max(v)))/(length(v)-1), where v is a vector of average expression values for different tissues.

### Cell Culture

ES-E14-Tg2a (E14) cells were grown on 0.1% gelatin-coated tissue culture dishes, in DMEM (Thermo Fischer, 41965-039) supplemented with 1x Non-Essential Amino Acids (Thermo Fischer 11140-035), 50 uM β-mercaptoethanol (Thermo Fischer 31350-10), 15% Fetal Bovine Serum (Thermo Fischer 10499-044), 500 Units/ml of Pennicilin/Streptomycin (Thermo Fischer, 15140122), and 100 Units/ml of Recombinant mouse LIF Protein (Merck ESG1107). Culture were seeded at an average density of ∼ 3.8*10^4^ cells/cm^2^ and passaged every 48 h.

### Analysis of cell cycle stage gene expression

We resuspended 10^6^ cells in 1 ml of PBS, containing 1 uL/mL of fluorescent reactive dye (LIVE/DEAD) and incubated in the dark for 30 minutes at room temperature. Cells were then washed with PBS, spun, and resuspended in fresh medium. We add 2 uL of DNA Hoechst 33324 dye (20 mM) and cells were incubated at 37°C for 20 minutes. Following incubation, cells were centrifuged at 4°C for 4 minutes at 400 g and the resulting cell pellet was resuspended in 500 ul of 3% FBS in PBS with 0.1% EDTA and kept on ice. Unstained and single dye controls prepared in parallel were used for FACS calibration.

Cells were sorted in an AstriosEQ (Beckmann Coulter) cell sorter and collected in 500 uL of RTL buffer from the RNAeasy extraction kit (QIAGEN). Forward and side scatter (FSC & SSC, 488nm DPSS laser) were used as is common for size and doublet exclusion. We excluded dead cells based on LIVE/DEAD fluorescence (488nm DPSS laser) and used DNA content (355nm DPSS laser) to define three gates: G1, S and G2/M (Supplementary Figure 5).

RNA was extracted using RNeasy mini kit (Qiagen) according to manufacturer’s instructions. RNA was reverse transcribed using Quatitect Reverse Transcription Kit (Qiagen 205313). Quantitative PCR analysis was performed using SYBR Green Quantitative PCR Master Mix (Roche 0692404001) in a Light Cycler 96 Real Time PCR system (Roche). Briefly, the RTqPCR reaction was assembled in 10uL with forward and reverse primers at a final concentration of .5 uM each, and SYBR Green Quantitative PCR Master Mix at a final concentration of 1X.

### Candidate lincRNA overexpression and cell cycle analysis

We used E-CRISPR [52] to design 2 guide RNA (gRNA) sequences located within 1 kb of lincCC1 and lincCC2 promoters respectively. We synthetized oligos containing these sequences and BbsI restriction sites. Annealed oligos were inserted into the pKLV-U6gRNA(BbsI)-PGKpuro2ABFP vector [53].

We seeded 1.5*10^5^ E14 cells/well in 6-well plates and allowed cells to grow overnight. We used lipofectamine 2000 (Thermo Fischer) to transfect 1000 ng of gRNA expression vector and 1000 ng of pAC95-pmax-dCas9-Vp160-2A-Neo [37] vector into mESC cultures overnight according to manufacturer’s instructions. We replaced growth medium 8 hours after transfection to minimize toxicity. Seventy-two hours post transfection one sample was collected for RNA to ensure the efficiency of the over-expression by qPCR; in parallel the other cells were pulsed with 10 uM Edu for 30 minutes in the dark at 37°C. Cells were washed with 1% BSA in PBS and trypsinized cells. Following a second wash with 1% BSA in PBS, cells were fixed at room temperature, protected from the light for 15 minutes in 100 uL of Click-iT fixative solution. After fixation, cells were washed in 1% BSA in PBS, resuspended in 100 uL of 1x Click-iT saponin-based permeabilization and wash reagent, and incubated at room temperature in the dark for 15 minutes. Following incubation, 500 uL of Click-iT reaction cocktail containing Alexa Fluor 488 Fluorescent Dye Azide were added and samples were incubated at room temperature in the dark for 30 minutes. Reaction was quenched by resuspending cells in 1 mL of 1x Click-iT saponin-based permeabilization reagent. To stain the samples for DNA content, 1 uL of Fx Cycle FarRed staining solution (Thermo Fischer F10348) was added to each sample, and Pure Link RNase A (Thermo Fischer 12091039) was added at a final concentration of 100 ng/mL. The samples were incubated at 4°C in the dark for 30 minutes, analyzed by flow cytometry analysis. Cells stained for DNA incorporation and content were analyzed using a 10 color/3 laser Beckman Coulter Gallios Analyzer. Blue (405 nm) and red (640 nm) excitation lasers were used for excitation of the Alexa Fluor 488 Fluorescent Dye Azide and Fx FarCycle Red respectively. Emission was detected using channels FL-09 and FL-06 for Alexa Fluor 488 and Fx FarCycle Red respectively. Three independent experiments were carried out for both candidate lincRNAs with at least 3 biological replicates each. 30-60k events were flowed per sample, except for the first experiment where 15-30k events were flowed.

Flow data was analyzed in FlowJo (v10.2). Flow events were gated on Cells and Singlets, and spillover was compensated. Low incorporation 2n and 4n populations, as well as a medium-to-high incorporation population, were easily identifiable in the DNA incorporation vs content scatterplot, and gated respectively as G1, G2/M and S (and ungated). This stage-level gating was performed by two gaters, blind to the direction and amplitude of target lincRNA expression modulation. Stage population fractions were comparable between gaters (Pearson correlations between stage fractions 0.65-0.97 depending on stage & experiment), and thus all results presented here are those of one gater.

Sample-level summaries of the flow cytometry results were generated using FlowJo (v10.2). Summaries were imported into R. Samples were quality-controlled based on their percentages of cells/singlets (indicative of cell health), and MFIs (indicative of staining strength). Experiments for which the associated qPCRs did not validate over-expression of the targets were excluded. Ultimately, 26 samples from 2 experiments and 5 treatment groups (scrambled, two different guides for lincCC1 and two for lincCC2) were retained. Differences in the percentages of cells in each stage were assessed between mock and treatment conditions using unpaired unequal variance t-tests.

## Acknowledgments

pKLV-U6gRNA-EF(BbsI)-PGKpuro2ABFP was a gift from Hiroshi Ochiai (Addgene plasmid # 62348; http://n2t.net/addgene:62348; RRID:Addgene_62348), pAC95-pmax-dCas9VP160-2A-neo was a gift from Rudolf Jaenisch (Addgene plasmid # 48227; http://n2t.net/addgene:48227; RRID:Addgene_48227).

We would like to thank the University of Lausanne Flow Cytometry Platform with help with FACS and sorting analysis. The computations were performed at the Vital-IT (https://www.vital-it.ch) Center for high-performance computing of the SIB Swiss Institute of Bioinformatics. We would like to thank Dario Bottinelli for technical support during the early stages of this project and Vincent Dion and Constance Ciaudo for reading and commenting on early versions of this manuscript. This work was funded by the Swiss National Science Foundation (grant PP00P3_150667 to A.C.M) and the NCCR in RNA & Disease (A.C.M.).

## Authors Contributions

AATS, AB and ACM contributed to the study design. AATS and JYT perfomed the insilico analysis. AATS, AB, MFS, BA and ACM performed the *in vitro* analysis. ACM supervised the study. ACM and AATS wrote the manuscript. All coauthors read and approved the manuscript.

## Figure legends

**Supplementary Figure 1:**

**A)** Schematic representation of the differential expression analysis pipeline. **B)** Scatterplot showing for each sample (cell) the total hit (mapped read) counts & “Cell Detection Rate”, i.e. fraction of genes detected by at least 1 hit in each cell. Low library quality exclusion thresholds are indicated by dashed magenta lines. Cells are coloured by cell cycle stage. **C)** Scatterplot showing for each cell the percent hits to mitochondrial (MT) genes & to ERCC spike-ins. Low library quality thresholds are indicated by dashed magenta lines. **D)** Distributions of Buettner’s cell size ratio [20] for cells in each stage. Magenta lines show medians +/-2 median absolute deviations; cells falling outside of these windows are excluded. **E)** Technical noise filter [44] as implemented in scLVM [20]: each gene is represented by its cross-sample mean expression (x-axis) and coefficient of variation (squared: CV2, y-axis). Blue dots correspond to ERCC spike-ins, while the green line represents the non-linear CV2 ∼ mean model that was fit to the spike-ins, and then used to test which genes show significantly higher-than-technical variability. Genes in black do not pass the significance test, those in red do. Counts of genes passing the test are indicated in the top-right hand corner.

**Supplementary Figure 2:**

**A)** boxplots of cross-sample mean expression (normalised shifted log10 counts) for genes of various classes: all=tested for differential expression. DE=differentially-expressed. NDE: non-differentially expressed. mRNA: protein-coding genes. lincRNA: lincRNA loci. **B)** Boxplots of shifted log10 mean loci expression for protein-coding genes (mRNA, dark grey) and lincRNAs (light grey), in both the sub-sampled (1/14th) data (SS) and FULL data from cell cycle staged single-cell RNAseq dataset [20].

**Supplementary Figure 3:**

Fold difference in relative expression for 9 predicted cell-cycle differentially expressed lincRNAs.

**Supplementary Figure 4:**

Genome browser view of the genomic location (mm10) of **A)** lincCC1 and **B)** lincCC2. Blue bars indicate conservation in multiZ alignments (60 vertebrates). **C)** gating strategy to identify cells from distinct cell cycle stages. Panel 1 shows the size exclusion gate based on FSC-H & SSC-H for removing debris. Panels 2 shows the FSC-H vs FSC-A doublet exclusion gate; a similar gate was implemented for SSC-H vs SSC-A. Panel 3 shows the partitioning of cells into 3 stages based on DNA content (measured on FL6-A) and EdU incorporation (measured on FL1-A).

**Supplementary Figure 5:**

Histogram of cellular DNA content measures as stained by DAPI, showing gates for delineating clearly-defined G1, S or G2/M stages.

