## Supplementary figures and images for "The contribution of lincRNAs at the interface between cell cycle regulation and cell state maintenance"

### Supplementary Figure 1

#### Supplementary Figure 1

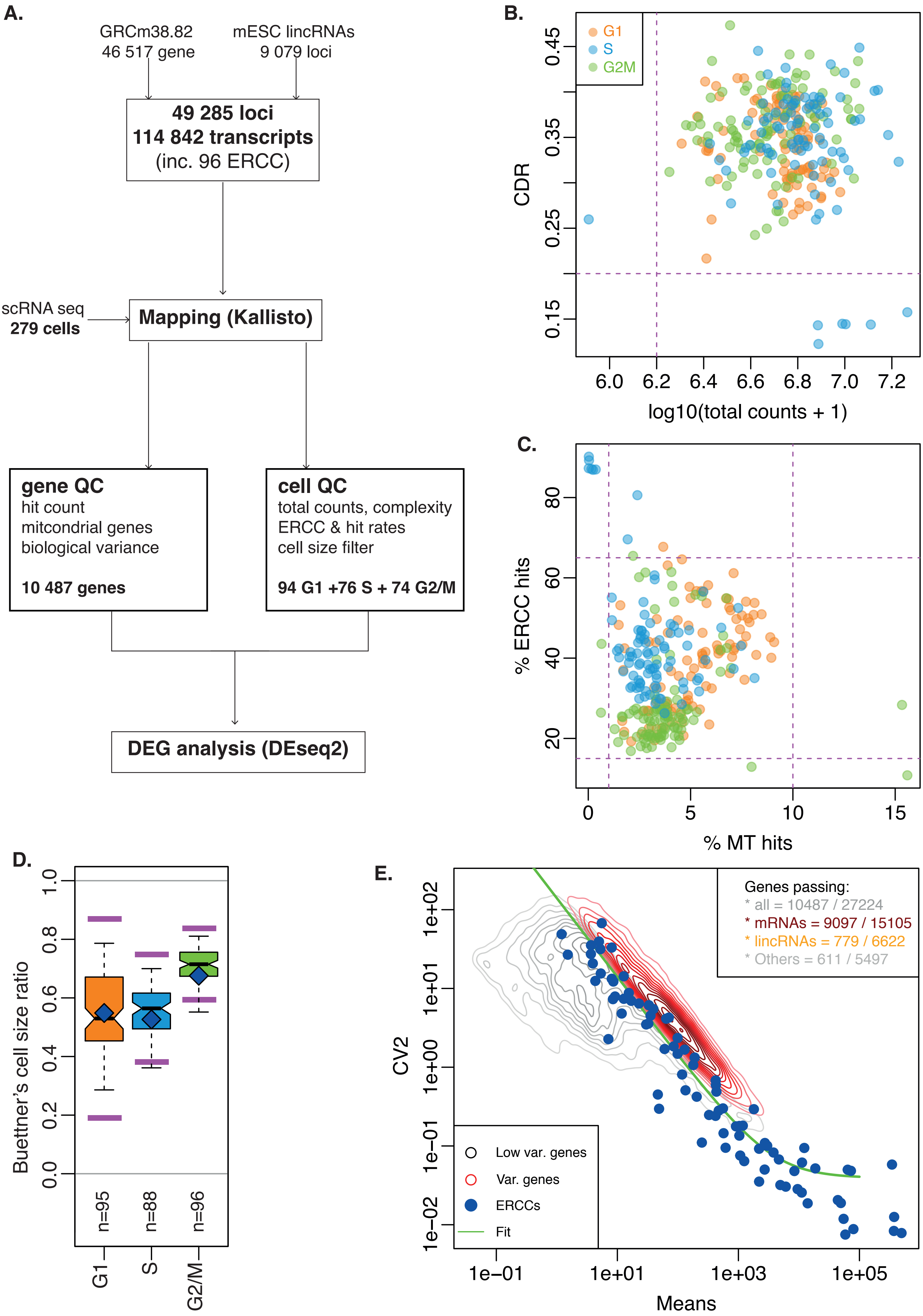

### Supplementary Figure 2

Supplementary Figure 2

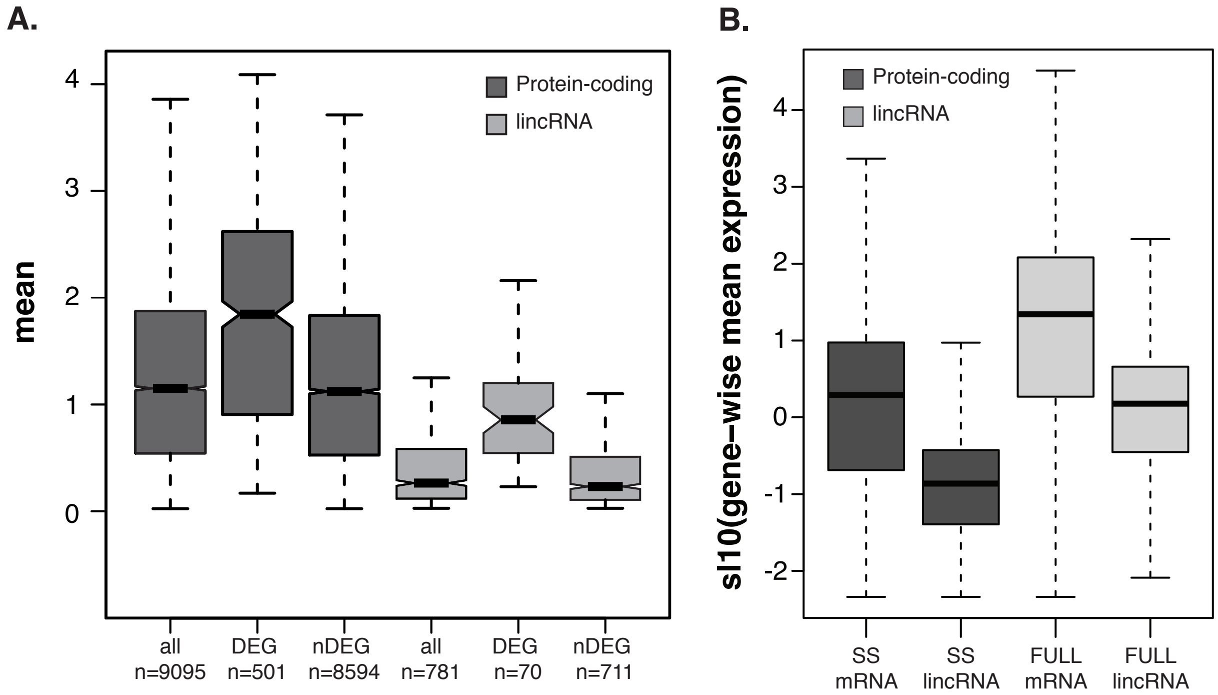


(Supplementary Figure 2).

### Supplementary Figure 3

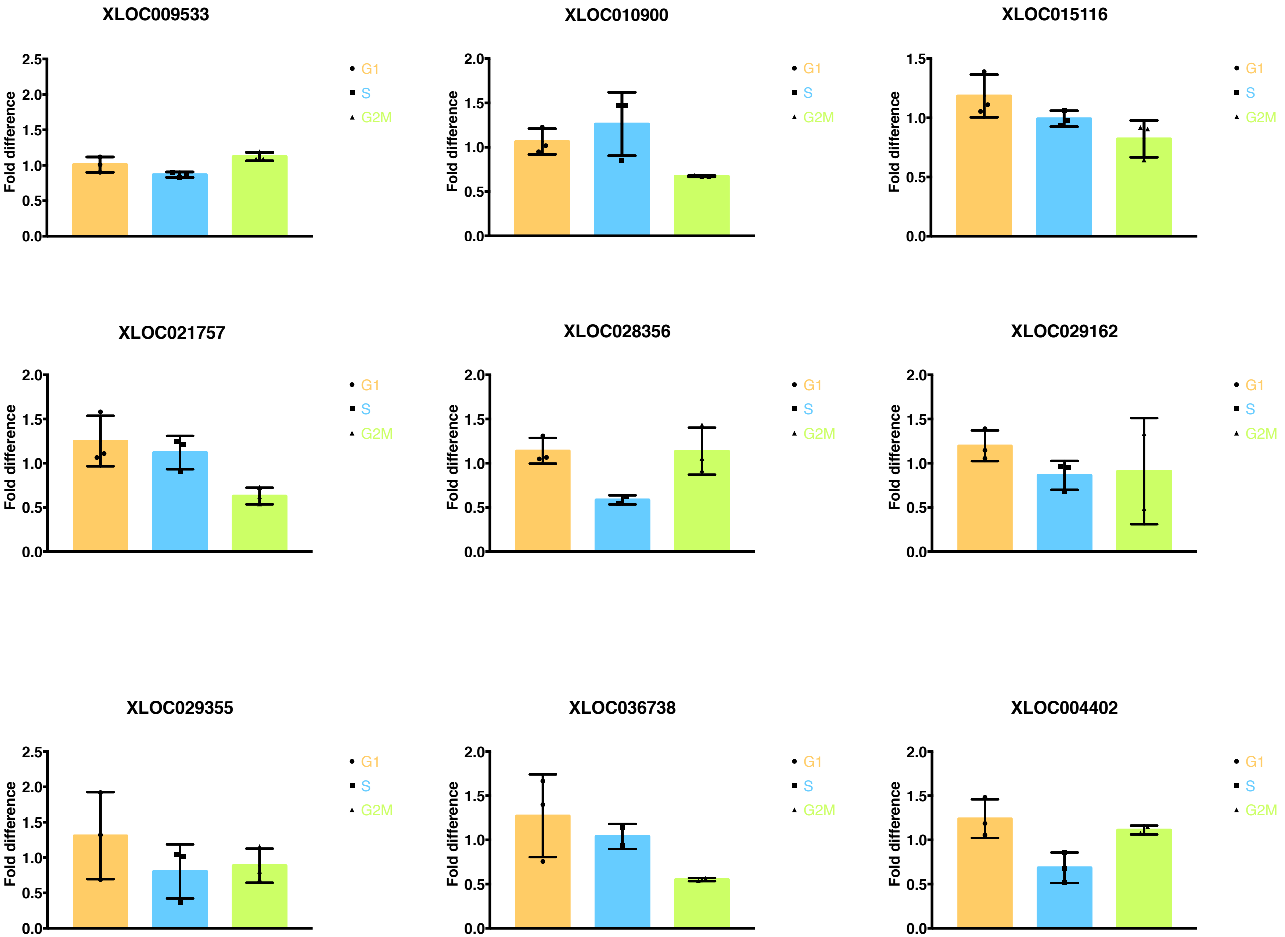

### Supplementary Figure 4

Supplementary Figure 4

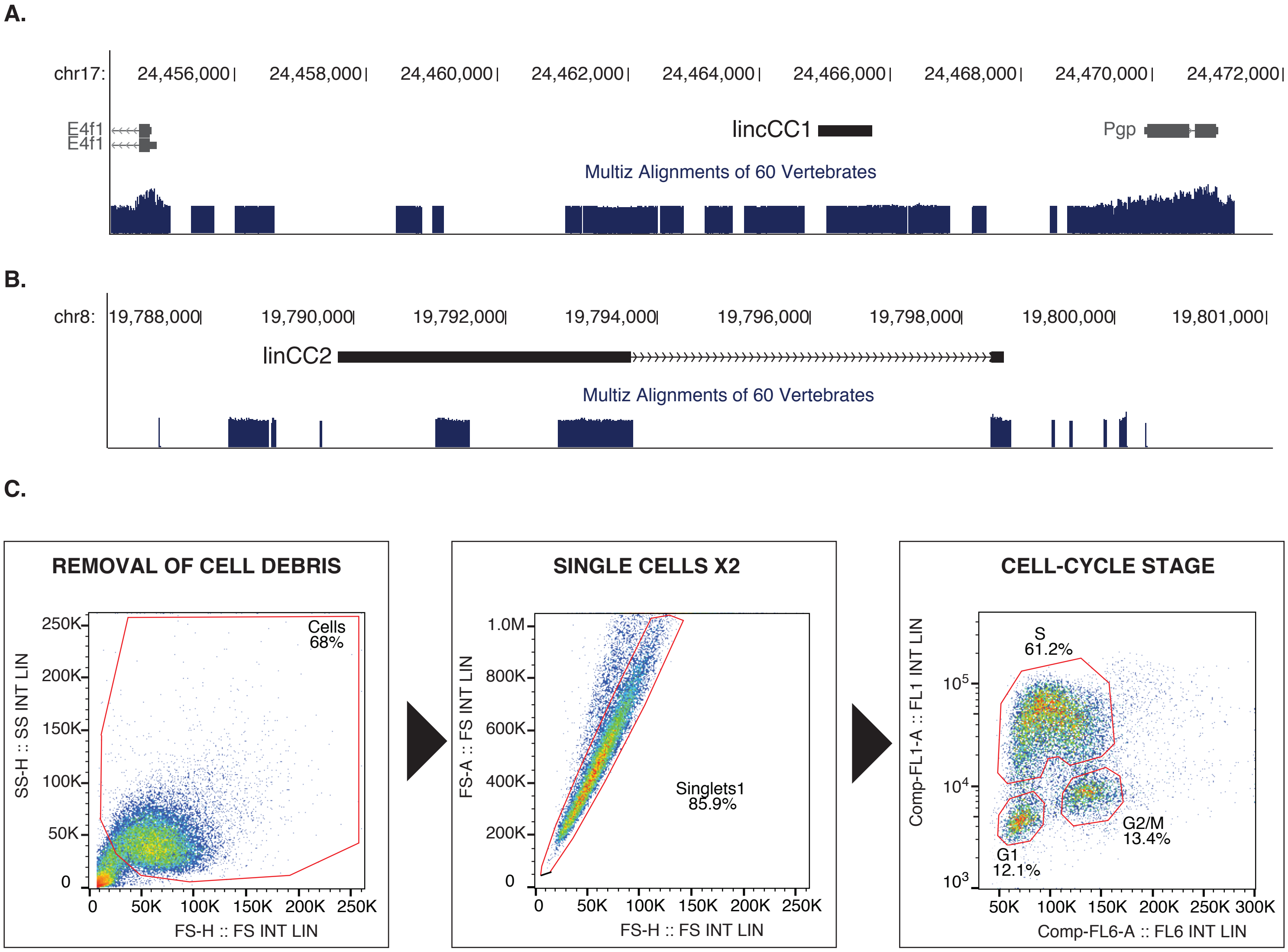

### Supplementary Figure 5

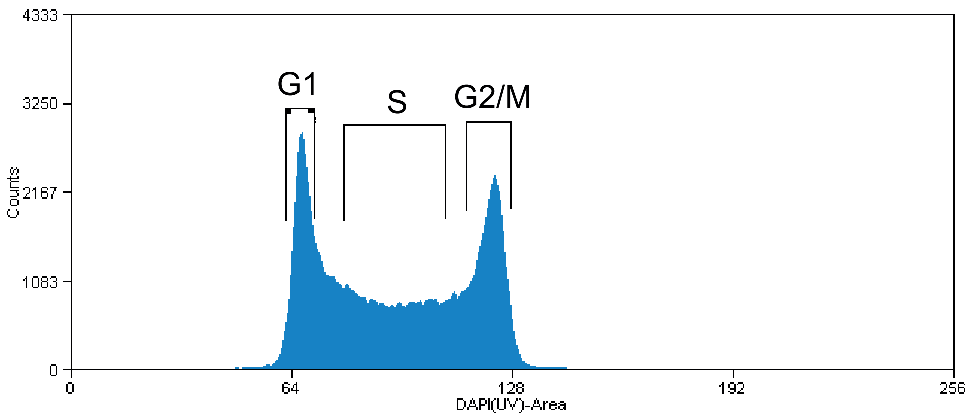
